# Chimeric single α-helical domains as rigid fusion protein connections for protein nanotechnology and structural biology

**DOI:** 10.1101/2020.09.29.318410

**Authors:** Gabriella Collu, Tobias Bierig, Anna-Sophia Krebs, Sylvain Engilberge, Niveditha Varma, Ramon Guixà-González, Xavier Deupi, Vincent Olieric, Emiliya Poghosyan, Roger M. Benoit

## Abstract

Chimeric fusion proteins are essential tools for protein nanotechnology. Non-optimized protein-protein connections are usually flexible, which makes them unsuitable as structural building blocks. Here we show that the ER/K motif, a single α-helical domain (SAH)^1^, can be seamlessly fused^2^ to terminal helices of proteins, forming an extended and partially free-standing rigid helix. Through the intrinsic stability of the SAH, two domains can be connected with a defined distance and orientation. We designed three constructs termed YFPnano, T4Lnano, and MoStoNano, and we show that a single SAH allows the connection of two separate structural domains with sufficient rigidity to form ordered crystals. The analysis of experimentally determined structures and molecular dynamics simulations reveals a certain degree of plasticity in the connections that allows the adaptation to crystal contact opportunities. Our data show that SAHs can be stably integrated into designed structural elements, enabling new possibilities for protein nanotechnology, for example to improve the exposure of epitopes on nanoparticles (structural vaccinology), to engineer crystal contacts with minimal impact in construct flexibility (for the study of protein dynamics), and to design novel biomaterials.

## Main text

The design of biomolecular architectures requires building blocks that can be integrated into larger assemblies in a predictable manner. While the prediction of nucleic acid folding and assembly in 2D and in 3D is very well established (e.g. Ref.^3^), the *de novo* design of proteins remains challenging despite recent advances in the field (summarized in Ref.^4^). Flexible, rigid and cleavable linkers are important components of recombinant proteins (reviewed in Ref.^5^). However, they often require extensive optimization to achieve the desired rigidity between the connected domains, unless a stable secondary structure of the linker - domain transition can be predicted, as in chimeric helices (reviewed in Ref.^2^).

Here, we explored the aptitude of single α-helical domains (SAHs), particularly an ER/K helix, as a modular component in extended chimeric helices. The stability of most α-helices depends on contacts with other regions in the tertiary structure of proteins. SAHs, on the other hand, are stabilized by dynamic charged interactions between the side chains that result in intrinsic stability and rigidity^1, 6^. SAHs can for example be found in smooth muscle caldesmon^7^, in myosin-X^8^, and in the ribosomal protein L9 from *Bacillus stearothermophilus^9^.* SAHs have been extensively characterized in solution using a wide variety of methods, including NMR^10, 11^, FRET^12^, and small angle X-ray scattering^13^, molecular dynamics simulations^8^, and they have been used as protein-protein spacers^14^. The stability of SAHs has furthermore been confirmed by crystallographic studies of such helices in their natural context (e.g. Refs.^8, 9^). The aim of our investigation is to test whether it is possible to rationally design rigid chimeric fusion proteins that are connected by a segment of a naturally occurring SAH. Such a connection, when introduced as a seamless extension of an N- or C-terminal helix, or of a helix near an accessible loop, only contacts the protein of interest at a single site. Unlike in direct chimeric helix fusions between two proteins^2^, an SAH helix provides an additional spacer that separates the two connected proteins sufficiently to avoid direct contacts between them **(Supplementary Fig. 1 a)** without requiring chemical crosslinking.

To test whether it is possible to use an ER/K segment as a modular building block to rigidly connect two separate protein domains in a variety of contexts, we designed three different fusion protein constructs; YFPnano, T4Lnano and MoStoNano.

### Design and structural characterization of YFPnano

We used a segment of the ER/K motif from myosin-X^8^ to engineer a fusion protein containing a rigid linker. Upstream of the rigid ER/K motif, we fused a calmodulin-binding peptide (CBP) from skeletal muscle myosin light-chain kinase (MLCK), which binds the protein calmodulin (CaM) with very high affinity^15^. The unbound CBP is a random coil^16^ that adopts an α-helical structure upon binding CaM, which wraps almost completely around the peptide^17^. Because the CBP from sMLCK, unlike the ER/K motif, is not intrinsically helical and rigid^16, 18^, it can be used as a flexible tag or linker that can later be rigidified by the addition of CaM. In our construct, the CBP-ER/K module was fused to the N-terminus of enhanced yellow fluorescent protein (EYFP, GFP-10C)^19^, which was used as a test protein that can be replaced by any protein of interest. CaM was fused to the C-terminus of YFP via a flexible linker, to allow intramolecular binding to the N-terminal CBP. We designed a single YFPnano construct, without any optimization of the linkers, and solved the structure by X-ray crystallography **(Fig. 1 a-d and Supplementary Table 1)**. The architecture of the final structure looks largely as expected. The C-terminal CaM is bound to the N-terminal CBP. The ER/K helix forms an extended chimeric helix with the CBP and the short N-terminal helix of YFP. The linker between YFP and CaM is flexible and hence not defined in the structure. The undefined region spans more than 20 residues, which is sufficient to connect the last defined amino acids, separated by ~3 nm (dashed line in Fig. 1 a and b).

**Fig. 1).**
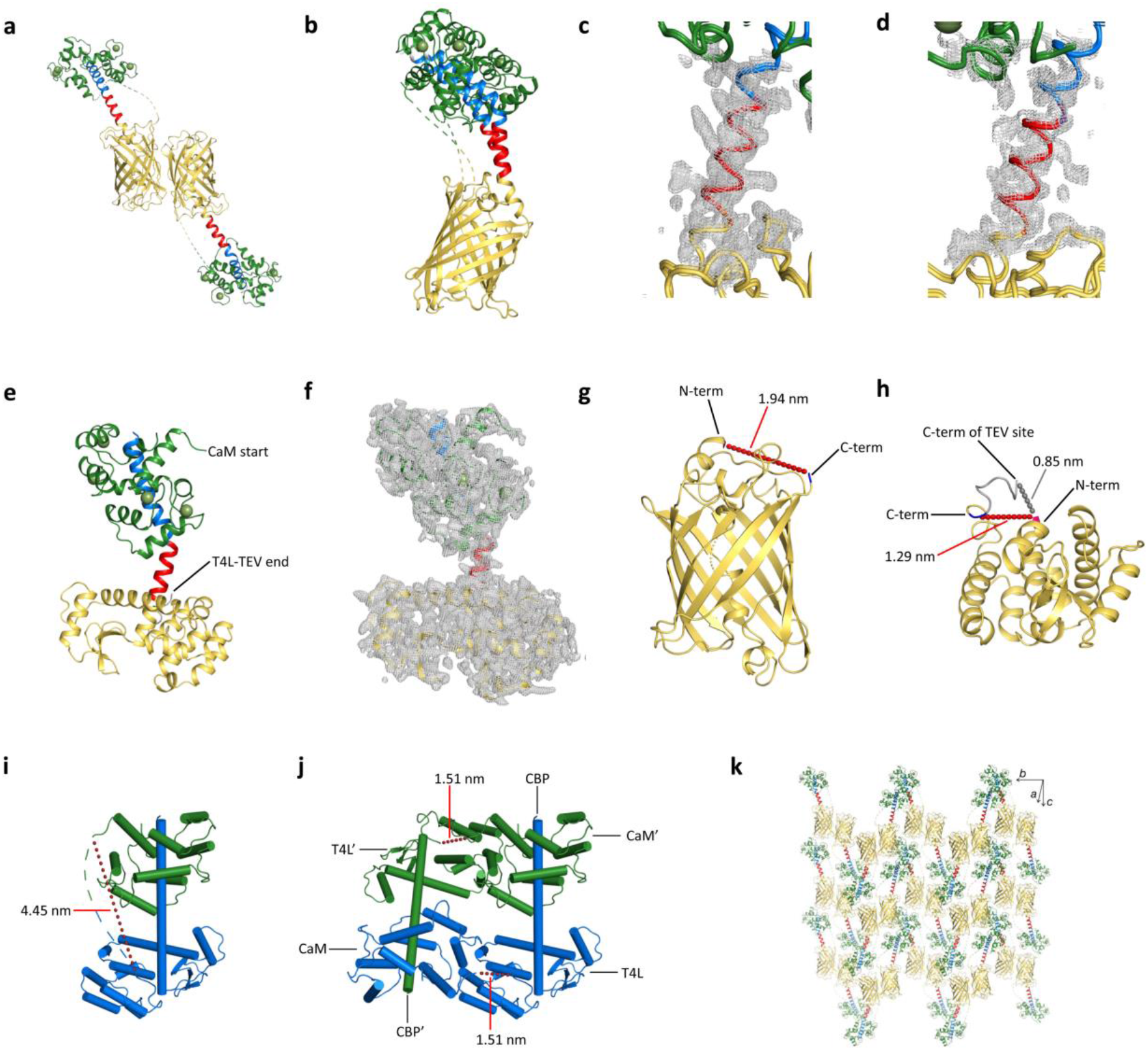
Crystal structures of the YFPnano and T4Lnano fusion proteins. **a-d,** Cartoon representations of the YFPnano structure: CBP (blue) - ER/K motif (red) - YFP (yellow) - linker (dashed lines) - CaM (green). Dashed lines indicate the flexible linker between YFP and CaM that is not visible in the structure. Calcium ions are shown as green spheres. **a,** The two molecules in the asymmetric unit (ASU) are shown. **b,** Superimposition of the YFP of chain B onto the YFP of chain A, emphasizing differences in the orientation of the CBP and ER/K helices. **c, d,** Electron density around YFPnano chains A (c) and B (d). 2Fo-Fc density map at a contour level of 1 σ displayed in a radius of 1.5 Å about each atom, showing that the core of the helix is well defined in the structure. The ER/K helix in chain A is stabilized by a crystal contact. The ER/K helix in chain B does not engage in direct crystal contacts and is fully exposed. The side chains are mostly undefined and B-factors are elevated in this region, however, there is clear electron density around the helix core, confirming that the ER/K motif indeed forms a stable and exposed helix. **e,** Cartoon representations of the T4Lnano structure: CBP (blue) - ER/K motif (red) - T4L (yellow) - linker (not shown) - CaM (green). **f,** 2Fo-Fc density map at a contour level of 1 σ displayed in a radius of 1.5 Å about each atom of the T4Lnano model. **g, h,** Geometry of the termini of YFP (g) and of T4L (h). In both proteins, the N- and C-terminus are in proximity to each other on the same side of the protein fold. **i, j,** Simplified cartoon representations of T4Lnano. Helices are shown as cylinders. **i,** The flexible linker that is not defined in the crystal structure (green/blue dashed line) is too short (9 residues, distance indicated with a red dashed line) to connect T4L with CaM within the same molecule. **j,** T4Lnano therefore has to form a dimer. The CaM of one T4Lnano molecule (blue) binds to the CBP of a symmetry-related molecule (green) in the crystal structure. The T4L-to-CaM distance is much shorter in the dimer (indicated with red dashed lines) and can be bridged by the undefined 9 residue linker. **k,** The crystal lattice of YFPnano, emphasizing the spacing introduced by the ER/K helix (red).

Analytical size-exclusion chromatography experiments confirm that YFPnano is monomeric **(Supplementary Fig. 2 a)**. There are two YFPnano molecules in the asymmetric unit, and in one molecule there is a kink at the transition between the CBP and the ER/K helix, probably induced by crystal packing forces **(Fig. 1 b)**. Ensemble-averaging FRET experiments using a Cerulean-ER/K-YFP fluorescent protein construct, support the role of the SAH as a spacer **(Supplementary Fig. 2 b and c)**.

### Design and structural characterization of T4Lnano

To test whether we can also apply this strategy to other proteins, we designed the fusion protein T4Lnano. Here, we kept the CBP - ER/K module and CaM from YFPnano but replaced the YFP by T4 lysozyme (T4L). The flexible linker connecting T4L to CaM is shorter than that connecting YFP to CaM in YFPnano **(Supplementary Fig. 3 a and b)**. Again, the SAH connection was sufficiently rigid to allow the formation of high-quality crystals without construct optimization. The resulting crystal structure is shown in **Fig. 1 e and f,** and the data collection and refinement statistics are listed in **Supplementary Table 1**.

The crystal structure of T4Lnano reveals that it forms a dimer. Unlike in YFPnano, only a short stretch (nine residues) of the flexible linker connecting T4L to CaM was undefined in the structure. This stretch is too short to span the 4.45 nm gap between the defined ends of the linker **(Fig. 1 i)**. On the other hand, the gap between the corresponding linker regions of T4Lnano and a symmetry-related molecule in the crystal lattice spans only 1.51 nm, which can be connected by a nine-residue linker **(Fig. 1 j)**. Thus, we conclude that in T4Lnano, the CaM - CBP binding is intermolecular, providing an example of a designed structural element that can form molecular assemblies.

### Molecular dynamics of T4Lnano

One of the two molecules in the asymmetric unit (ASU) of the YFPnano crystal structure shows a kink in the CBP near the transition to the ER/K motif. A similar kink is also present in the T4Lnano crystal structure, which contains one molecule in the ASU. To study whether this kink is induced by crystal packing forces, we performed classical all-atom molecular dynamics (MD) simulations of T4Lnano in solution, using the crystal structure as a starting point. We simulated a single CaM-bound CBP - ER/K - T4L entity (as shown in **Fig. 1 e and i)**, disregarding the flexible linker. First, we run ten independent replicas of 100 ns each (10 x 100 ns). As shown in **Figs. 2 a and b**, in the absence of crystal contacts, the kink between the CBP and ER/K helices tends to significantly straighten or completely disappear in most of the simulations.

**Fig. 2).**
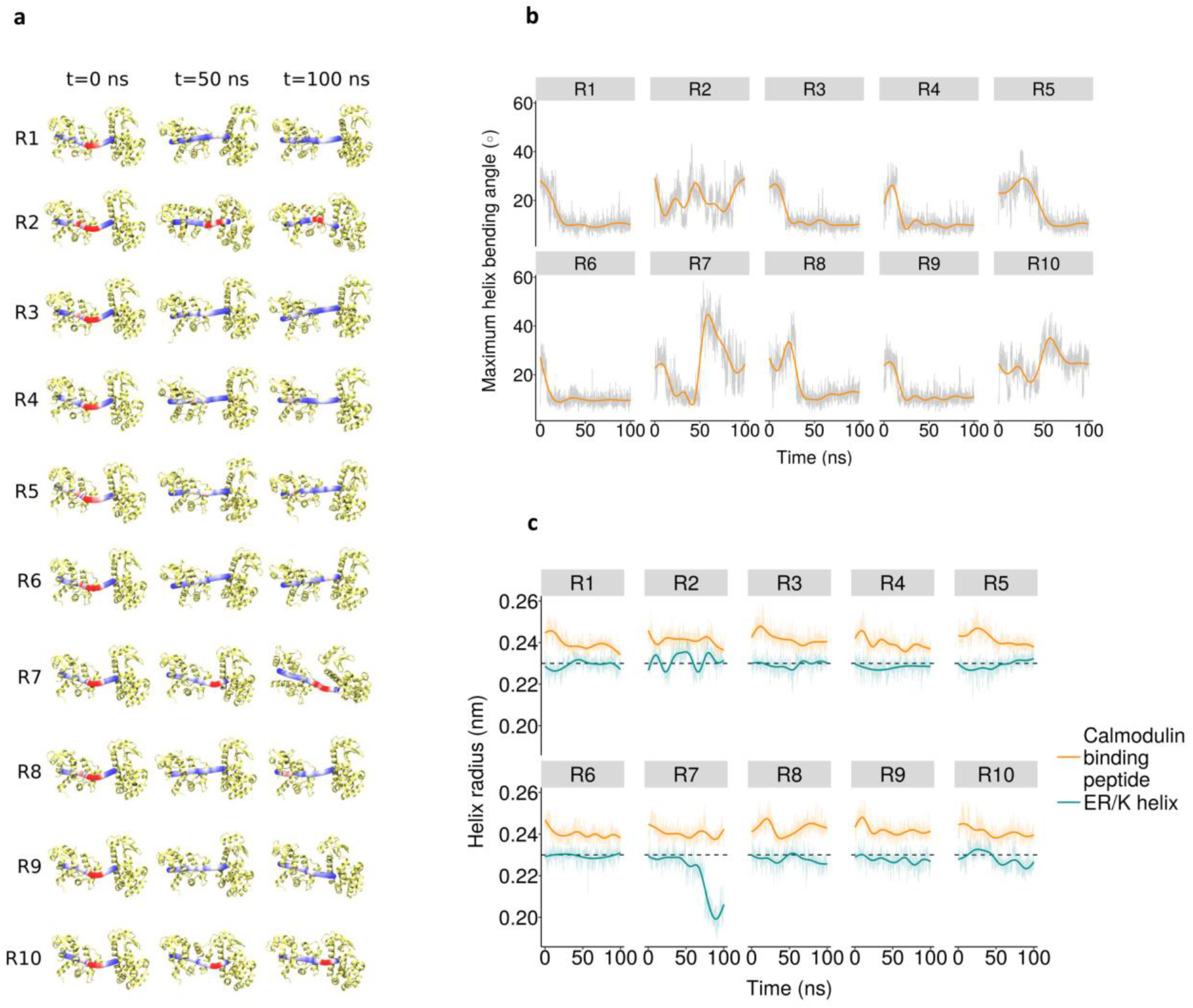
Molecular dynamics simulations. **a,** Helix bending angle across all simulations of T4Lnano. The figure shows three snapshots (at the beginning (0 ns), middle (50 ns), and end (100 ns) of the trajectory), for all 10 replicas. Bendix^20^ was used to calculate and visualize the curvature of the CBP-ER/K helix (residues 6 to 36). A gradient color scale, (blue (low) to red (high)), is used to represent maximum angle values of the helix during the simulation. The rest of the protein is depicted in light yellow cartoons. **b,** Time series plots of maximum helix bending angles. Maximum helix bending angles of the CBP- ER/K helix (i.e. residues 6 to 36) over time are depicted as light gray lines for every simulation replica of T4Lnano (10 x 100 ns). Overlaid bold orange lines represent smoothed averages. **c,** Time series plots of helix radius. Comparison of the helix radius over time between CBP (residues 6 to 25, yellow) and the ER/K helix (residues 26 to 36, blue). Bold lines are smoothed averages. A black dashed horizontal line shows the radius value of an ideal helix (0.23 nm).

**Supplementary Movie 1** illustrates this straightening, using replica 1 as a representative simulation. To further support these results, we extended the first three simulation replicas to 500 ns (i.e. 3 x 500 ns). As shown in **Supplementary Fig. 4**, the CBP-ER/K helix displays a similar straightening tendency at longer time scales. We also analyzed the simulation data to calculate the structural regularity of the CBP-ER/K domain by measuring the radius of each helix segment. We observe that the radius of the ER/K motif averaged over all simulations is 0.228 ± 0.005 nm, which corresponds to the ideal value for an α-helix of 0.23 nm (Ref.^21^). A time series plot **(Fig. 2 c)** shows that despite some fluctuations this value was maintained in almost all replicas during the trajectories, indicating that the ER/K helix has a very high propensity to remain helical in solution with minimal protein-protein contacts. The CBP helix also shows an average radius (0.241 nm ± 0.003 nm) corresponding still to an α-helix **(Fig. 2 c)**. In this case, the CBP is likely stabilized via interactions with surrounding protein domains.

### CBP-CaM interaction

Overall, the CaM-CBP binding mode in our YFPnano and T4Lnano structures is in good agreement with the solution NMR structure described by Ikura et al.^17^, based on which our constructs were designed. The CaM-CBP interaction is described in detail in the **Supplementary Text and Supplementary Fig. 5**. In summary, the calcium-sensing protein calmodulin consists of two domains. The N-terminal domain comprises amino acids 4 −74 and the C-terminal domain comprises residues 82-147. The sMLCK peptide (CBP) is buried in a large hydrophobic channel of CaM, the CBP-CaM interaction is dominated by hydrophobic interactions.

To ensure formation of a stable complex, we used a CBP sequence with a single-point amino acid mutation (Asn-to-Ala) in the MLCK M13 peptide that results in super-high affinity to CaM (K_d_ = 2.2(±1.4) × 10^−12^ M, corresponding to a ~1000-fold affinity improvement compared to the wt peptide)^15^. To the best of our knowledge, our YFPnano and T4Lnano structures are the first structures of CaM in complex with the M13 MLCK peptide containing this mutation.

The CaM-CBP interaction can be used for intramolecular domain bridging or for the connection of two separate protein domains^22^. In YFPnano, we included the CBP and the CaM within the same construct, to promote an intramolecular interaction. In T4Lnano, CBP and CaM were also included in the same construct, however with a shorter linker that prevented intramolecular binding, resulting in homodimer formation. When designing fusion proteins that contain both, CBP and CaM, it is of key importance to take into consideration the geometry of exposure of the CBP with respect to CaM. If intramolecular binding is geometrically hindered, constructs form dimers (e.g. T4Lnano) or oligomers with intermolecular CBP-CaM interactions. Such self-assembling “protein strings” can be interesting for other protein nanotechnology applications, for example for the design of novel biological materials **(Supplementary Fig. 1 b)**.

Strings and scaffolds with a wide range of properties can be designed, depending on amino acid composition and geometry. Chimeric SAH-containing proteins are modular building blocks of adjustable length and rigidity. The generation of tunable biomaterials for medicine, such as hydrogels^23^, and of peptide-based biopolymers^24^ for the production of biodegradable textiles, are two of many examples of possible applications.

### Design and structural characterization of MoStoNano

For our third construct, MoStoNano, we designed a fusion protein involving a more complex protein assembly. As one of the components, we chose the molybdenum storage protein (MoSto) from *A. vinelandii*^25^ which consists of a dimer of trimers. The trimers themselves consist of three heterodimers, each comprising an α-subunit and a β-subunit. The overall structure can hence be described as (αβ)_3_ (Ref.^26^). The β-subunit has an N-terminal α-helix that we extended into an SAH. To the N-terminus of the SAH-extension (ER/K segment from Myosin-X^8^), we fused a thioredoxin 1 (Trx-1) from *E. coli*^27^, a small globular soluble protein that ends in an α-helix. The structures of each individual component are shown in **Fig. 3 a**. In a first non-optimized construct, Trx-1 and the SAH were flexible and hence not visible in the structure (data not shown). Thus we designed two additional constructs **(Fig. 3 b)** with optimized ER/K - Mosβ transitions. The construct with the shortest transition yielded high-quality crystals and initial refinement steps after molecular replacement revealed electron density for the ER/K helix and Trx-1 **(Fig. 3 c)**.

**Fig. 3).**
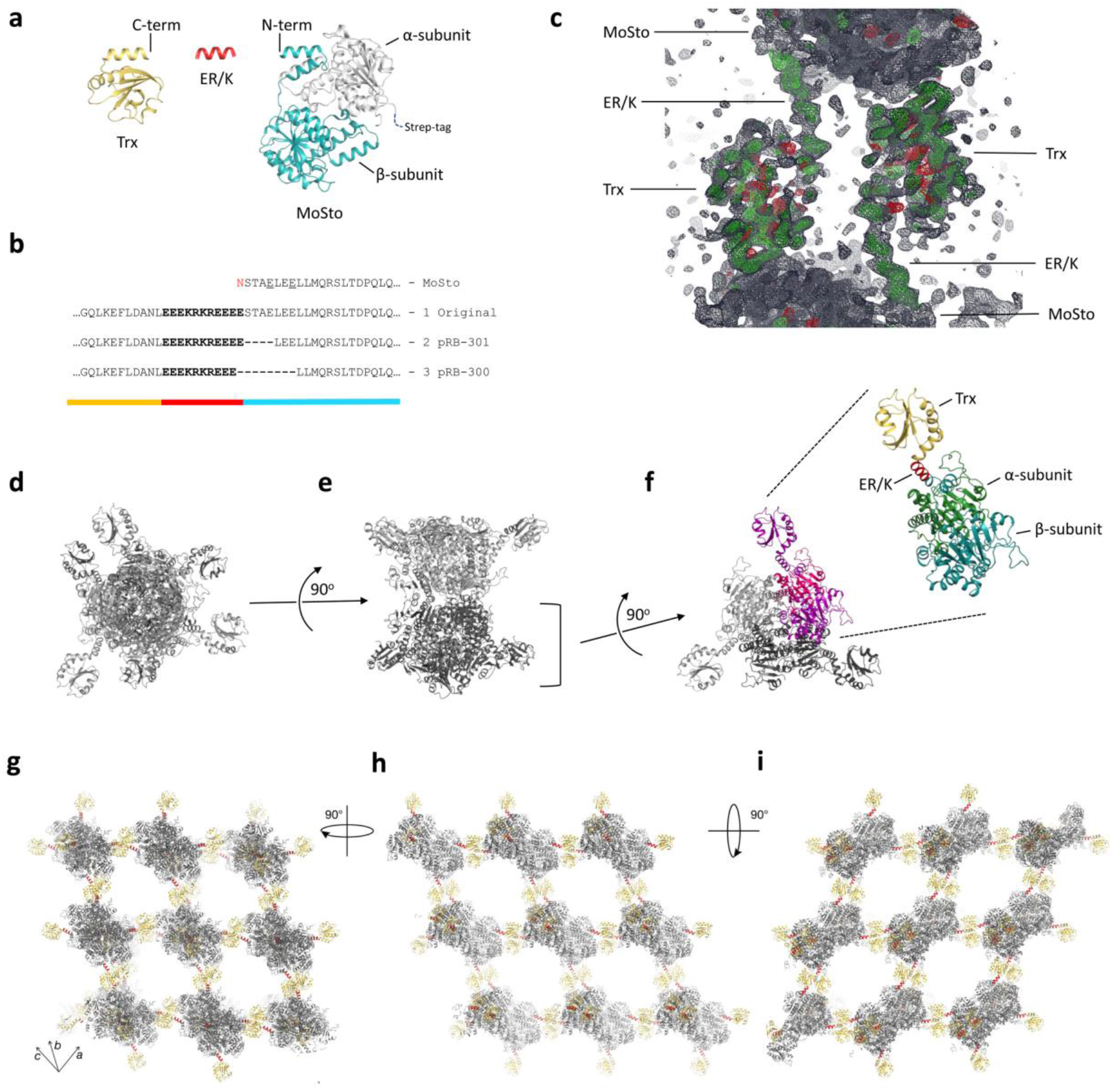
Crystal structure of the MoStoNano fusion protein. **a,** Components that were used to design MoStoNano: Thioredoxin 1 (yellow, PDB entry 3DYR, **Ref.^27^)**, a segment of the ER/K helix from myosin X (red, PDB entry 5HMO, **Ref.^8^**) and the β-subunit of a molybdenum storage protein (blue, PDB entry 4F6T, **Ref.^25^**). The α-subunit of the molybdenum storage protein (grey) was expressed as a separate chain with a C-terminal Strep-tag to isolate the complex. **b,** Sequences of the different constructs at the Trx-1 - ER/K - MoSto β-subunit transition. In the X-ray structure of construct 1, the Trx-1 and ER/K were not resolved. Construct 2 crystallized but the diffraction quality was low and did not allow structure elucidation. In construct 3 (= MoStoNano), electron density for Trx-1 and the ER/K was clearly apparent in the structure. **c,** Initial refinement after molecular replacement, using only the MoSto model, revealed density for Trx-1 and the ER/K helix. 2Fo-Fc density is shown in dark blue and Fo-Fc density is shown in green. **d,** Top view and **e,** Side view of the biological unit of MoStoNano. The heterodimer comprising one α- and one β-subunit of MoSto forms a trimer. This trimer then forms a dimer. In (d) and (e), one of the trimers is shown in light grey, the other in dark grey. **f,** Top view of one of the two trimers. One heterodimer is shown in purple (β-subunit, including the Trx-1 and ER/K) and pink (α-subunit). The other two heterodimers are shown in light and dark grey, respectively. The close-up view shows the detailed components of one MoStoNano heterodimer. The space group of the crystal was *H32* and the asymmetric unit contained one (αβ)-dimer. **g, h, i,** The crystal lattice of MoStoNano from three views, illustrating how each MoSto biological unit (grey) is embedded between Trx-1 - ER/K elements (yellow and red), that mediate crystal contacts to symmetry-related MoSto molecules. A segment of the ER/K helix is free-standing (see also (c) and (f)).

Crystal contacts are mediated through thioredoxin 1 - mosβ crystal contacts **(Fig 3 g-i)**. The ER/K helix has a free-standing and solvent-exposed region, that provides separation between the architectures of Trx-1 and MoSto. The large MoSto complex is embedded in the crystal lattice with only single-helix connections contributing to spacing and crystal stability in one region. This is reflected by high B-factors and limited definition of the electron density in the Trx-1 - ER/K region of the structure.

A closer look at the crystal lattices of MoStoNano **(Fig. 3 g-i)** and YFPnano **(Fig. 1 k)** reveals another striking characteristic of the seamless ER/K fusions: through the spacing introduced by the SAH, it is possible to promote the formation of crystal contacts at a significant distance from a protein region of interest, while retaining conformational freedom in the latter within the crystal lattice, which is ideal for time-resolved crystallography **(Supplementary Fig. 1 c)**.

### Structure elucidation of MoStoNano by cryo-EM

We then evaluated whether the ER/K connection in MoStoNano is sufficiently rigid for the display of small proteins on a large scaffold for structural characterization by cryo-EM **(Fig. 4 a and b)**. Initial attempts were limited by the presence of preferential orientations of the particles on the cryo grids (data not shown). After extensive optimization, the protein retained only a slight tendency for preferential orientations **(Fig. 4 c)**. We therefore elucidated the cryo-EM structure with imposed D3 symmetry, resulting in an overall resolution of 2.9 Å **(Fig. 4 b and d)**, but the electron densities of Trx-1 and the ER/K helix were missing in the 3D classification and refinements, irrespective of the imposed symmetry, possibly due to the helix plasticity observed in our crystallography data and molecular dynamics calculations.

**Fig. 4).**
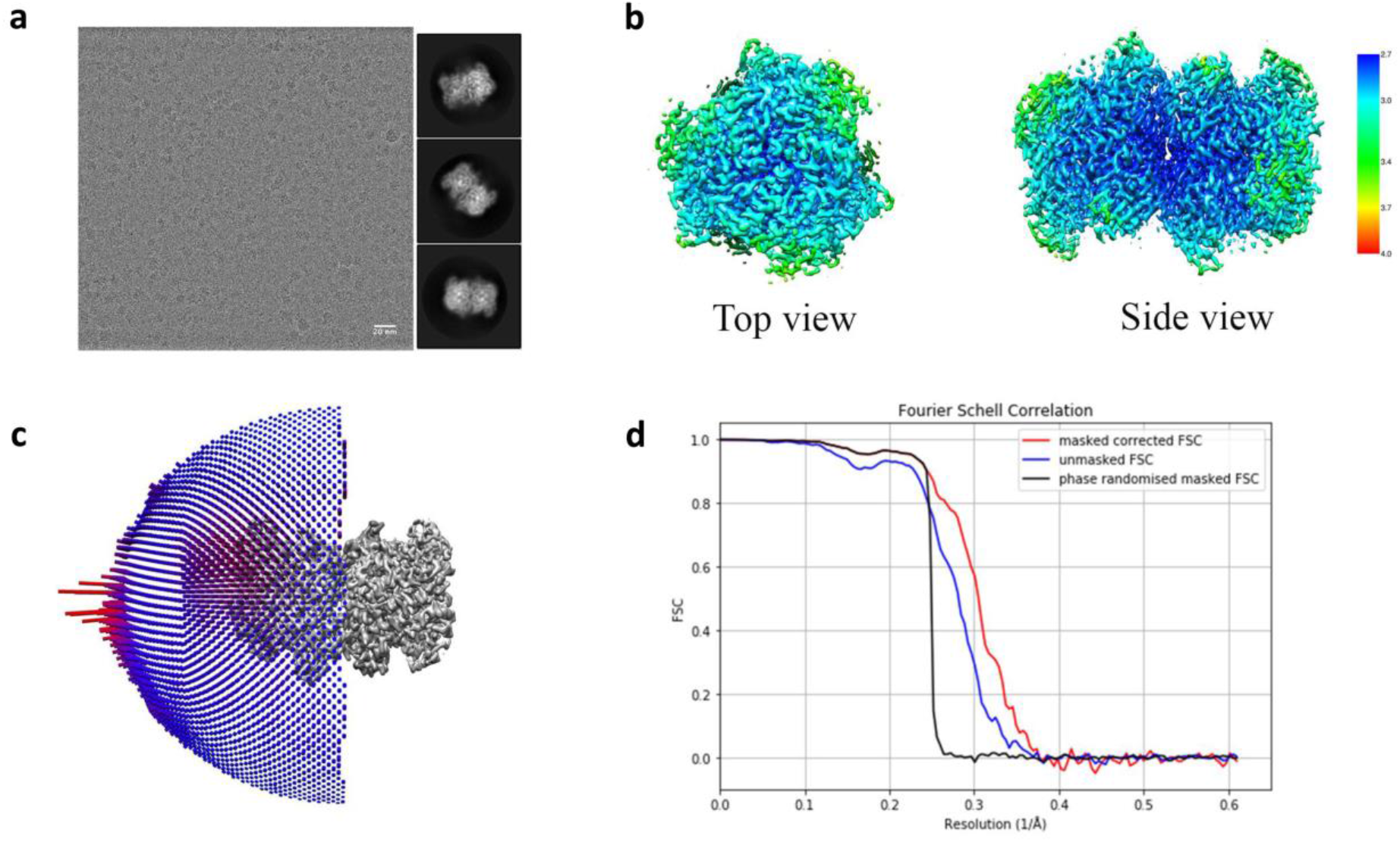
Cryo-EM structure refinement and local resolution of MoStoNano. **a,** Left: Representative micrograph after applying the beam induced motion correction procedure, scale bar 20 nm. Right: Representative 2D class averages used for the final reconstruction with a box size of 24.6 nm. **b,** local resolution maps with representative top and side views of the reconstruction of the protein with imposed D3 symmetry. The resolution is color coded as indicated by the scale bar. **c,** Distribution of Euler angles of all the particles that were used for the final 3D reconstruction. **d,** Fourier Shell Correlation curves of the 3D reconstruction of MoStoNano with imposed D3 symmetry from the RELION auto-refine procedure.

A homogeneous protein sample is typically assumed to be a prerequisite for crystal formation. The observation that our SAH-fusion proteins readily crystallized suggests that, despite some degree of structural fluctuations, the fusion proteins in solution had preferred conformations that were sufficiently predominant to provide building blocks that integrated into the growing crystals. Such connections with prevalent conformations can be of great significance to improve the accessibility of protein or peptide epitopes displayed on nanoparticles **(Supplementary Fig. 1 d)**. Interestingly, some important advances in vaccinology have been based on structure-based improvement of antigen presentation^28^.

Together, our data show that it is possible to rationally design seamless and extended protein helices containing a free-standing ER/K segment, for their use as modular structural elements that connect two protein domains with limited flexibility. This protein nanotechnology approach, based on molecular biomimetics, is highly useful for a wide range of applications, such as the improvement of the exposure of epitopes on nanoparticles (structural vaccinology), the engineering of crystal contacts with a minimal impact on construct flexibility (for the study of protein dynamics), and the design of novel biomaterials. In principle, our approach allows to connect any protein containing a C-terminal α-helix to any protein containing an N-terminal α-helix, and insertions within proteins (e.g. at helix-loop transitions) are also possible. The use of circular permutation^29^ to provide a terminal helix can increase the repertoire of amenable proteins.

## Materials and Methods

### Cloning

### YFPnano

The cDNA encoding calmodulin (UniProt P0DP23) was amplified by PCR using the primers 5’-AACTTCAAGGATCCGCTGCTGCTGCTGACCAGCTGACCGAAGAAC-3’ and 5’-TGCTCGAGTGCGGCCGC TCATTATTTAGCGGTCATCATCTGAACGAACTC-3’. The PCR product was digested with BamHI and NotI and ligated into the BamHI / NotI sites of a modified pET vector containing eYFP. The eYFP-TEVsite-linker-CaM region of the resulting 6xHistag-PreScissionSite-eYFP-TEVsite-linker-calmodulin construct was amplified with the primers 5’-CGCGAAGAGGAAGAAGTGAGCAAGGGCGAGGAGCTGTTC-3’ and 5’-CTCGAATTCGGATCCTCATTATTTAGCGGTCATCATCTGAACGAACTCTTCGTAG-3’ and subcloned into a modified, PCR-linearized^30^ pET vector, containing segments coding for an N-terminal calmodulin-binding peptide^15^ and a rigid ER/K helix^8^, by co-transformation cloning^31, 32^. Plasmid linearization was performed using primers 5’-TTCTTCCTCTTCGCGTTTGCGCTTTTCCTC-3’ and 5’-GGATCCGAATTCGAGCTCCGTCGACAA-3’.

### T4Lnano

A FragmentGene comprising an N-terminal His-tag, a PreScission cleavage site, the calmodulin-binding peptide, the ER/K motif, T4 lysozyme, a flexible linker containing a TEV cleavage site, and calmodulin was ordered from Genewiz (codon optimized for expression in *E. coli*) and cloned into the NcoI and EcoRI sites of pET-28a (Novagen).

### MoStoNano

A bicistronic FragmentGENE coding for both subunits of molybdenum storage protein (MoSto) from *Azotobacter vinelandii*^26^, with a segment coding for thioredoxin 1 and an ER/K helix fused in frame to the N-terminus of the mosβ subunit, and a C-terminal, TEV-cleavable Strep-tag in the mosα subunit, was ordered from Genewiz, codon optimized for expression in *E. coli.* The gene fragment was cloned into the NcoI and EcoRI sites of a Novagen pET-28a vector. The regions flanking the ER/K helix were then modified through plasmid linearization by PCR, followed by blunt end ligation.

### FRET constructs

His-tag - PreScissionSite - mCerulean - (linker) - eYFP - TEVsite - TwinStrepTag constructs were ordered from Genewiz, codon optimized for expression in *E. coli,* cloned in the NcoI and NotI sites of pET-28a (Novagen). The linker region consisted of the sequence EEEKRKREEEE in the SAH construct. The ER/K - to - eYFP transition amino acid sequence was identical to YFPnano. In the GS-linker construct, the amino acid sequence GGGGSGGGGSG replaced the ER/K segment. In the construct without a linker, mCerulean was directly fused to eYFP.

### Protein expression and purification

#### YFPnano and T4Lnano

The fusion proteins were expressed in *E. coli* cells (Rosetta2(DE3) for YFPnano and BL21(DE3) (EMD Millipore Novagen) for T4Lnano, in 2x YT medium supplemented with 50 μg/ml kanamycin. The cells were grown at 37 °C, 130 rpm in a shaker-incubator until an OD_600_ of 0.75 was reached. CaCl_2_ was added to a concentration of 0.2 mM, and overexpression of the fusion proteins was induced by the addition of IPTG to 1 mM. Expression was carried out overnight at 20 °C. The cells were then harvested by centrifugation. The cell pellets were resuspended in 50 mM Tris, pH 7.8, 150 mM NaCl, supplemented with a tablet of complete EDTA-free protease inhibitor cocktail (Roche Diagnostics). The cells were lysed by ultrasonication. After incubation with lysozyme and DNase I at 4 °C for 30 minutes, the lysates were clarified by centrifugation at 16500 x g for 15 minutes at 4 °C. ß-mercaptoethanol was added to a final concentration of 5 mM to the clarified lysates. The proteins were purified by Ni-NTA IMAC using Ni-Sepharose 6 Fast Flow resin (GE Healthcare). The wash buffer consisted of 50 mM Tris, pH 7.8, 150 mM NaCl and the elution buffer of 50 mM Tris, pH 7.8, 150 mM NaCl, 350 mM imidazole. After elution, DTT was added to a concentration of 5 mM. To cleave off the 6xHis-tag, recombinant human rhinovirus 3C protease was added to the eluted proteins, and the proteins were dialyzed overnight against wash buffer. The cleaved fusion proteins were further purified by gel filtration on a Superdex 75 column in 20 mM Tris, pH 7.8, 150 mM NaCl (YFPnano) or 25 mM Tris, pH 7.5, 150 mM NaCl (T4Lnano). CaCl_2_ was added to the eluted proteins to a concentration of 5 mM.

### MoStoNano

The protein was expressed in BL21(DE3) *E. coli* cells in LB (Luria-Bertani) medium supplemented with 50 μg/ml kanamycin. The cells were grown at 37°C, 180 rpm in a shaker incubator until an OD_600_ of 0.6 was reached. Overexpression of the fusion protein was induced by the addition of 0.5 mM IPTG and the temperature was lowered to 32°C for overnight expression. Cells were harvested at 4°C and the pellet was resuspended in 50 mM TRIS pH 8.0, 1 M NaCl, 5% glycerol, 1 mM MgCl_2_, 1 mM Na_2_MoO_4_, 1 mM ATP, 1 mM DTT, 0.2% Tween20, supplemented with complete EDTA-free protease inhibitor cocktail (Roche Diagnostics) and DNAse I (Roche Diagnostics). Lysis was performed at 4°C using Emulsiflex C3 (Avestin) at 1500 PSI in 4 cycles.

The lysate was clarified by centrifugation at 43000 rpm for 45 minutes at 4°C. The supernatant was applied to a StrepTrap HP column (GE Healthcare) for purification. The wash buffer consisted of 50 mM TRIS pH 8.0, 10 mM NaCl, 5% glycerol, 1 mM MgCl_2_, 1 mM Na_2_MoO_4_, 1 mM ATP, 1 mM DTT and 0.2% Tween20. The elution was performed using 50 mM TRIS pH 8.0, 10 mM NaCl, 5 mM D-desthiobiotin, 5% glycerol, 1 mM MgCl_2_, 1 mM Na_2_MoO_4_, 1 mM ATP, 1 mM DTT and 0.2%Tween20.

The protein was further purified by size exclusion chromatography, using a Superdex200 Increase column (GE Healthcare) in 50 mM MOPS pH 6.5 and 50 mM NaCl.

### FRET constructs

The fusion proteins were expressed in BL21(DE3) *E. coli* cells at 20 °C overnight, using IPTG induction (1 mM). The proteins were purified by Ni-NTA IMAC using Ni-Sepharose 6 Fast Flow resin (GE Healthcare). The wash buffer consisted of 50 mM Tris, pH 7.8, 150 mM NaCl and the elution buffer of 50 mM Tris, pH 7.8, 150 mM NaCl, 350 mM imidazole. The proteins were further purified by gel filtration on a Superdex 75 column in 50 mM Tris, pH 7.5, 150 mM NaCl, 5 mM CaCl_2_.

### Crystallization and structure elucidation

#### YFPnano

For crystallization, the YFPnano fusion protein was concentrated to 20 mg/ml in a 10 kDa cutoff centrifugal filter device (Amicon). Initial crystals were obtained in the Qiagen PEGs Suite crystallization screen, well H2 (0.2 M di-ammonium tartrate, 20% (w/v) PEG 3350). Optimized crystals were grown in 24% (w/v) PEG 3350, 0.2 M di-ammonium tartrate, 10% glycerol in sitting drops at 20 °C. The cryobuffer consisted of reservoir solution including 25% ethylene glycol.

A complete dataset to 1.9 Å resolution was collected from a single crystal at 100 K at the X06DA beamline of the Swiss Light Source synchrotron at the Paul Scherrer Institute in Villigen-PSI, Switzerland, using a wavelength of 1.0 Å. XDS^33^ was used for data processing. The high-resolution limit was chosen according to the CC1/2^34^, making sure that the *I*/σ*I* in the highest resolution shell was above 1. The space group of the crystal was *P21.* The structure of the fusion protein was solved by molecular replacement using a structure of YFP (PDB entry 3V3D^35^) as the search model. Two YFP molecules could be positioned in the asymmetric unit using Phaser^36^. After the first refinement cycles, additional density for the extended helix and calmodulin became apparent. A helix was placed into the density near the N-terminus of YFP chain A in Coot^37^. In iterative rounds of refinement and model building, the helix was built, starting at the very characteristic electron density of tryptophan 4 and phenylalanine 17. Calmodulin was then placed by first building the most characteristic helix in Coot^37^, followed by superposing the calmodulin structure from PDB entry 2BBM^17^ onto the CBP helix and the CaM helix. After initial refinement, CaM regions that did not fit the electron density were deleted and then manually built in Coot^37^ with iterative rounds of refinement and model building. The same strategy was then applied to build the extended helix and CaM in chain B. Refinement was performed using phenix.refine^38^. The geometry and stereochemistry was validated using MolProbity^39^. The final refined structure exhibits good geometry and stereochemistry. In chain B, the side chains of CaM amino acids 41-72 are poorly defined and amino acids 73 - 79 are undefined. The flexible linker connecting YFP and CaM is undefined in both chains.

### T4Lnano

The T4Lnano fusion protein was concentrated to 36 mg/ml in a 10 kDa cutoff centrifugal filter device (Amicon). Initial crystals were obtained in the Qiagen PEGs II Suite crystallization screen, well H5 (8% PEG 8000, 200 mM LiCl, 50 mM MgSO_4_). Optimized crystals were grown in 8% PEG 8000, 200 mM LiCl, 100 mM Tris pH 8.0, 15% glycerol in sitting drops at 20 °C. Reservoir solution including 30% glycerol was used as the cryobuffer.

The structure was solved by sulphur-SAD at the beamline X06DA at the Swiss Light Source, Villigen-PSI, Switzerland. Nineteen 360° data sets were collected on two different crystals at a wavelength of 2.07 Å in various *chi* settings of the PRIGo goniometer^40, 41^. With 0.2° oscillation and 0.1 s exposure per frame, this corresponds to a dose of ~ 0.5 MGy per data set. The data were processed using XDS and merged using XSCALE^33^. The space group was *P*2_1_2_1_2. *CRANK2*^42^ was used for substructure determination, phasing and initial model building. An additional high-resolution data set to 2.1 Å resolution was collected on a third crystal using a wavelength of 1.0 Å **(Supplementary Table 1)**. XDS^33^ was used for data processing. After sulphur phasing, T4L from PDB entry 1LYD^43^ and the calmodulin and calmodulin-binding peptide from our YFPnano structure (Chain A) were superimposed onto the initial model and then used for molecular replacement with the high resolution dataset. The high resolution model was built in iterative rounds of refinement and model building using Coot^37^ and phenix.refine^38^. The geometry and stereochemistry was validated using MolProbity^39^. The final refined structure exhibits good geometry and stereochemistry. In the loop comprising amino acids 287-292, the side chains are undefined. The flexible linker connecting T4L and CaM is partially undefined.

### MoStoNano

The protein for crystallization was in 50 mM MOPS pH 6.5, 50 mM NaCl and was concentrated to 3.8 mg/ml. Crystals formed in the Qiagen NeXtal PEGs II Suite screen, well E3 (0.1 M tri-Sodium citrate pH 5.6, 10% PEG 4000, 10% Isopropanol). 25% of glycerol were used as the cryoprotectant. A complete dataset to 2.85 Å resolution was collected from a single crystal at 100 K at the X06DA beamline of the Swiss Light Source synchrotron in Villigen-PSI, Switzerland, using a wavelength of 1.0 Å.

The autoPROC pipeline was used to process the data, diffraction frames were integrated using XDS^33^, the integrated intensities were scaled with AIMLESS^44^ and POINTLESS^45^ and an anisotropy correction was performed automatically with STARANISO^46^. The space group of the crystal was *H*32. The structure of the fusion protein was solved by molecular replacement using a structure of MoSto (PDB entry 4F6T, Ref^25^) as the search model. One dimer consisting of one α- and one β-subunit of MoSto could be positioned in the asymmetric unit using Phaser^36^. After initial refinement cycles, clear difference density for thioredoxin 1 and the ER/K helix became apparent. Two helices were placed into the thioredoxin density in Coot. Thioredoxin 1 from *E. coli*^27^ (PDB entry 3DYR) was then superimposed onto the helices and the ER/K helix connecting thioredoxin and mosB was manually build in Coot^37^.

This initial model was then optimized through iterative rounds of refinement and model building using respectively BUSTER^47^ and COOT. Translation–liberation–screw-rotation provided in the BUSTER program were applied at each refinement cycle. Atomic coordinates and structure factor amplitudes have been deposited in the Protein Data Bank after validation with MolProbity. PyMOL^48^ and Coot^37^ were used for the preparation of the figures of the crystal structures. UCSF Chimera^49^ was used for the preparation of the cryo-EM structure figures.

### FRET experiments

The Förster resonance energy transfer (FRET) experiments were performed using an excitation λ of 434 nm and an emission λ of 529 nm for FRET, on a Biotek Synergy 4 plate reader (90% gain). The protein concentration was 1mg/ml and the sample volume was 50 μl. Each sample was provided in triplicates in a 384 well plate.

### MD simulations

The T4Lnano crystal structure was first pre-processed using VMD1.9.3^50^ to remove any co-crystallization atoms (other than water molecules) closer than 5 Å to the protein. The structure was then placed into a water box and the system was neutralized with ions while adjusting the ionic strength using the CHARMM-GUI builder^51^. Except for Glu28, Glu35, Glu327 and His66, all titratable residues of the protein were left in the dominant protonation state at pH 7.0. No mutations or addition/deletions of residues were made to the native sequence of the protein. Prior to the production runs, the geometry of the system was optimized by energy minimization and further relaxed by a sequence of equilibration steps. In these equilibration steps, harmonic positional restraints were applied to all C_α_ atoms of the protein and gradually released throughout the equilibration run. Then, ten independent trajectories were spawned from the last snapshot of the equilibrated system using a random seed. Production simulations for each replica were run in the NPT ensemble at 1,013 bar and 303,15 K. All simulations were run using gromacs v2020^52^ in combination with the CHARMM36m force field^53^. Gromacs v2020^52^, Bendix^20^, and VMD1.9.3^50^, were used to post-process and analyze all trajectories. Figures were prepared using VMD1.9.3^50^, and the R ggplot2 library^54^.

### Preparation of the cryo-EM grids

The protein peak obtained from the size exclusion chromatography was collected and diluted to 0.4 mg/ml. Cryo-EM grids were prepared by applying 3.5 μl of protein to the glow-discharged UltrAufoil R1.2/1.3 200-mesh grids. The grids were blotted for 3.5 s, plunge-frozen in liquid ethane using Vitrobot Mark IV operated at 4°C and 100% humidity (Thermo Fischer Scientific) and stored in liquid nitrogen until the day of cryo-EM data collection.

### Cryo-EM data collection and processing

Initial screening of the grids was performed using a JEM2200FS (JEOL) microscope with an in-column Omega energy filter, operated at 200 kV and equipped with a K2 Summit direct electron detector (Gatan). Final data were collected on a Titan Krios (Thermo Fisher Scientific), equipped with a Gatan Quantum-LS Energy Filter and a Gatan K2 Summit direct electron detector at C-CINA in Basel. The microscope was operated at 300 kV using SerialEM software^55^.

Data were collected in a counting mode with dose fractionation and a total dose of ~62 e^-^/Å^2^ at a magnification corresponding to a calibrated pixel size of 0.82 Å. A total of 3,400 movies were collected in 2 days of data collection. The dataset was pre-processed and pruned in FOCUS^56^ using MotionCor2^57^ with dose weighting for movie alignment, and CTFFIND4^58^ for micrograph contrast transfer function estimation. A total of 2,472 movies were selected based on iciness, CTF resolution, and defocus values for further data processing.

Particles were first manually picked in RELION 3.0^59^ from the resulted 2,472 CTF corrected micrographs and subsequently auto-picked using a template generated from the manual picking procedure, which resulted in 199,441 particles. After several cycles of 2D classification, the best classes were selected, resulting in ~96,000 particles for initial 3D classification, for which we used a wild-type structure of MoSto as an initial mode (EMDB 4907), which had been low-pass filtered to 60 Å to avoid any reference bias. After 3D classification, particles were 3D refined, CTF refined and post-processed, which resulted in a map at a resolution of 2.9 Å. Local resolution was determined using RESMAP 1.1.4^60^ with filtering produced using RELION 3.0.

## Acknowledgements

We thank May Sharpe, Laura Vera, Meitian Wang and the MX group for support at the Swiss Light Source (SLS) beamlines and at the crystallization facility. We also thank Martin Schärer, Tobias Weinert, Tim Grüne and Andrea Prota for important discussions about structural biology related aspects of the project, and Takashi Ishikawa, Gebhard F.X. Schertler, Jan Pieter Abrahams, Michel Steinmetz and Dmitry Veprintsev for supporting the project. We thank Steffen Brünle for providing important information about MoSto expression, Timothy Sharpe for biophysics advice, and Hans Widmer, Sandra Jacob and Gregor Cicchetti for interesting discussions about the project. We thank Mohamed Chami and Lubomir Kovacik at C-CINA Basel for support in cryo-EM data collection, and the EM Facility at the PSI for technical support in cryo-EM data collection. This work was supported by grants from Novartis FreeNovation, UBS Promedica (1401/M) and the Swiss National Science Foundation (SNF SPARK, CRSK-3_190414) to R.M.B., and a grant from the Swiss National Science Foundation (192780) to X.D..

## Author contributions

The project was initiated and coordinated by R.M.B. The fusion proteins were designed by R.M.B. and cloned by T.B., G.C., N.V. and R.M.B.. The proteins were expressed and purified by G.C., T.B., A.S.K. and N.V. and crystallized by A.S.K., T.B., G.C. and R.M.B.. Sulfur phasing of T4Lnano was performed by S.E., V.O. and R.M.B.. The crystal structures were solved and refined by R.M.B., S.E., T.B. and V.O.. The FRET experiments and the analytical SEC experiments were performed by T.B.. Molecular dynamics simulations were carried out and analyzed by R.G.-G. and X.D.. Grids for cryo-EM data collection were prepared by G.C. and screened by E.P. and G.C., the cryo-EM structure was elucidated by E.P.. The manuscript was written by R.M.B. with contributions from all authors.

## Data availability

Structure coordinates have been deposited in the PDB database under accessions codes 6HR1 (YFPnano), 6XYR (T4Lnano) and 6YT3 (MoStoNano).

## Supplementary Material

**Supplementary Fig. 1).**
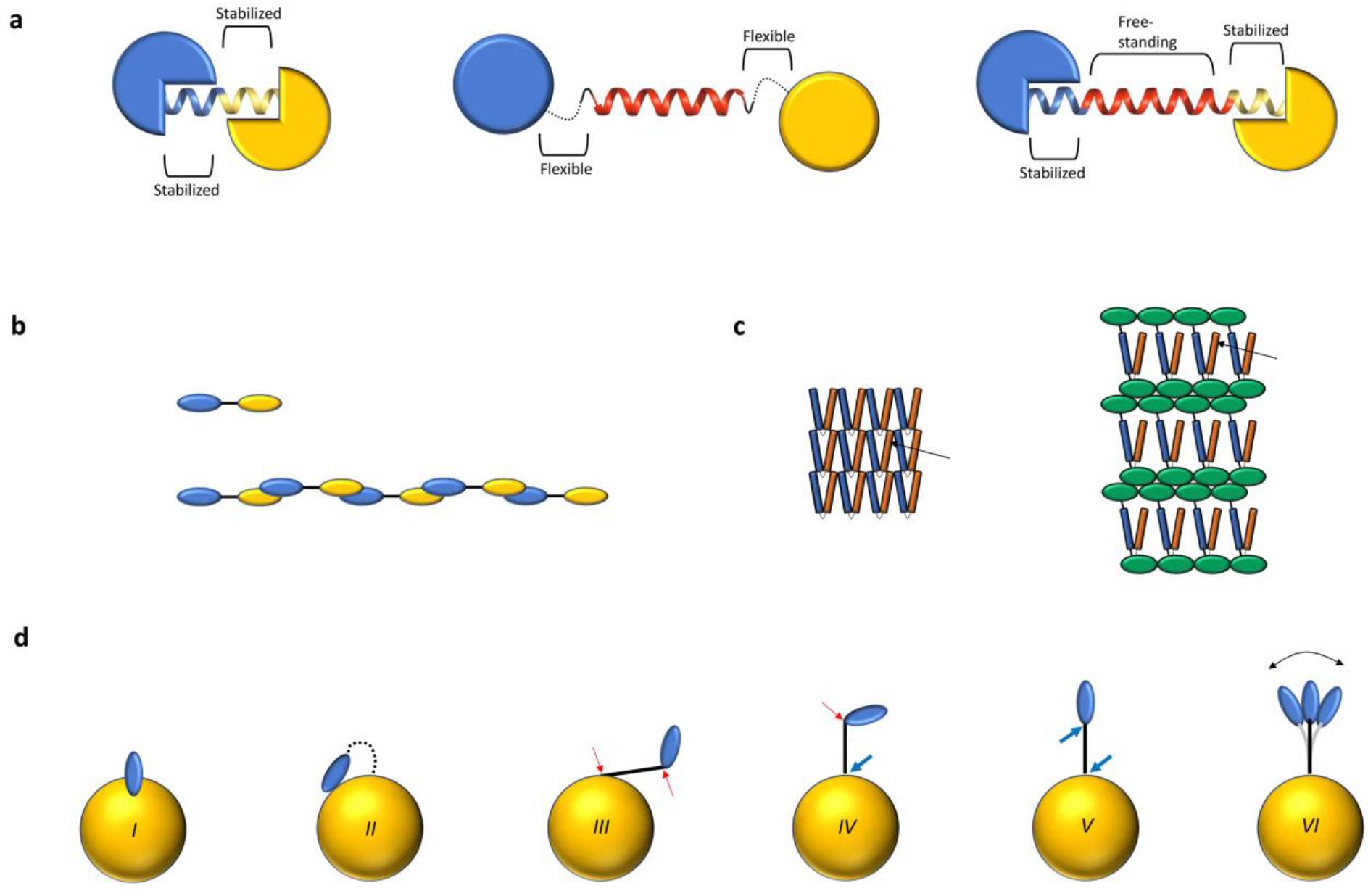
Rigid chimeric helices. **a, Left:** Direct helix fusion (e.g.^2, 32, 61, 62^). The terminal helices are part of the fused proteins (blue and yellow) and are stabilized by tertiary structure, with no or only a short free-standing region. **Middle:** Use of an SAH (red helix) as a spacer (e.g. Ref.^14^). The connections to the proteins are not optimized for rigidity. **Right:** Rigid, extended SAH-containing chimeric helix, as shown here in YFPnano, T4Lnano and MoStoNano. The connected terminal helices of the proteins form an extended, intact helix containing an intrinsically stable, free-standing SAH segment (shown in red). **b-d,** Potential applications for rigid chimeric helices. **b,** Schematic example where two domains (blue and yellow) with a strong binding affinity to each other, connected by a rigid SAH (black line), can oligomerize into a designed molecular string. **c,** Engineering of crystal lattices to optimize the conformational freedom of a protein region within the crystal lattice, schematic representation. **Left:** Crystal lattice of a protein (comprising one blue and one yellow helix, connected by a loop shown as a dotted line). The arrow indicates the region of interest, which cannot move freely due to crystal contact clashes. **Right:** Engineered protein containing a fusion protein (green) that is inserted into the protein of interest through one rigid linker (solid black line) and one flexible linker (dotted line). Here, the region of interest (yellow helix, arrow) can freely move within the crystal lattice. **d,** Nanoparticles for the presentation of epitopes. The multimeric protein nanoparticle is shown schematically as a yellow sphere. The viral epitope (protein or peptide) is depicted in blue. The linker is shown as a solid black line (rigid) or as a dotted black line (flexible). For clarity, only one of many linker-epitope entities is shown.***I,*** Fusion without linker. Parts of the epitope can be inaccessible due to steric hindrance. ***II,*** If a flexible linker is used, the epitope may fold back and interact with the nanoparticle, again resulting in partial inaccessibility. ***III,*** a rigid linker, used as a spacer, can improve the display, even if the attachment points to the nanoparticle and to the epitope allow hinge movements. Flexible hinges are indicated by red arrows. ***IV*** and ***V***, we here suggest the engineering of rigid connections (bold blue arrows) to further improve epitope presentation. ***IV,*** Using one rigid connection and one flexible hinge, the epitope is displayed like a flag that can move freely. ***V*,** Using two rigid connections, the epitope is displayed in a specific orientation. ***VI,*** Our molecular dynamics experiments and structures suggest that the rigid chimeric SAH fusions retain some conformational freedom, with a strong tendency towards a straight, helical conformation. The importance of designed self-assembling nanoparticles as improved vaccines has been summarized in Ref.^28^: Since the origins of vaccination, empiric development of vaccines has allowed us to prevail against most diseases. However, the development of vaccines against some diseases, such as the respiratory syncytial virus (RSV), remained difficult until recently. Through multiple steps of structure-based engineering^63, 64^, an effective vaccine has been developed by *in silico* design. The self-assembling nanoparticle is stable in a conformation with ideal antigen presentation, resulting in optimal immunogenicity and adjuvanticity^28^.

**Supplementary Fig. 2).**
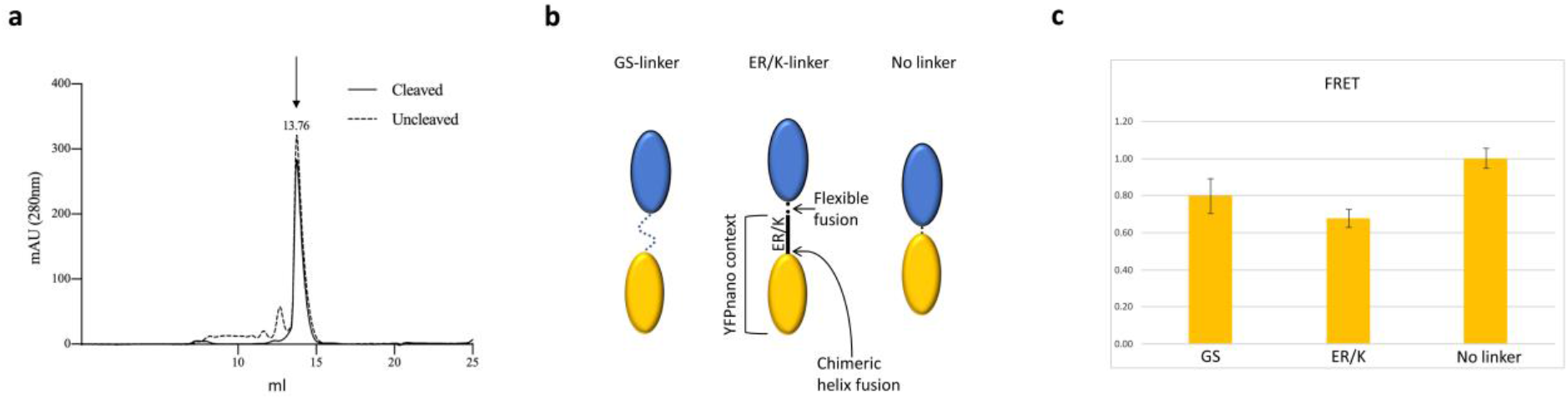
Size-exclusion chromatography (SEC) and Förster resonance energy transfer (FRET) experiments. **a,** YFPnano contains a TEV protease cleavage site in the flexible linker between the C-terminus of YFP and the N-terminus of CaM. The main size-exclusion chromatography (SEC) peak, indicated by the arrow, elutes at the same retention volume with the uncleaved and the cleaved protein, indicating that CaM-to-CBP binding is intramolecular in YFPnano. **b,** Schematic representation of the three constructs used for FRET experiments**. Left:** An mCerulean fluorescent protein was fused to an eYFP fluorescent protein via a flexible Glycine-Serine (GS) linker. **Middle:** In the second construct, the linker consisted of a seamless ER/K motif - eYFP fusion (identical to YFPnano). To the N-terminal side of the ER/K helix, mCerulean was fused via a short flexible connection. **Right:** In the third construct, the linker was omitted. **c,** Sensitized-acceptor emission ensemble averaging FRET measurements of the three constructs. Although the GS linker can extend more than the helical ER/K linker, less energy transfer was observed for the ER/K construct, indicating that the ER/K segment acts as a spacer, separating mCerulean and eYFP.

**Supplementary Fig. 3).**
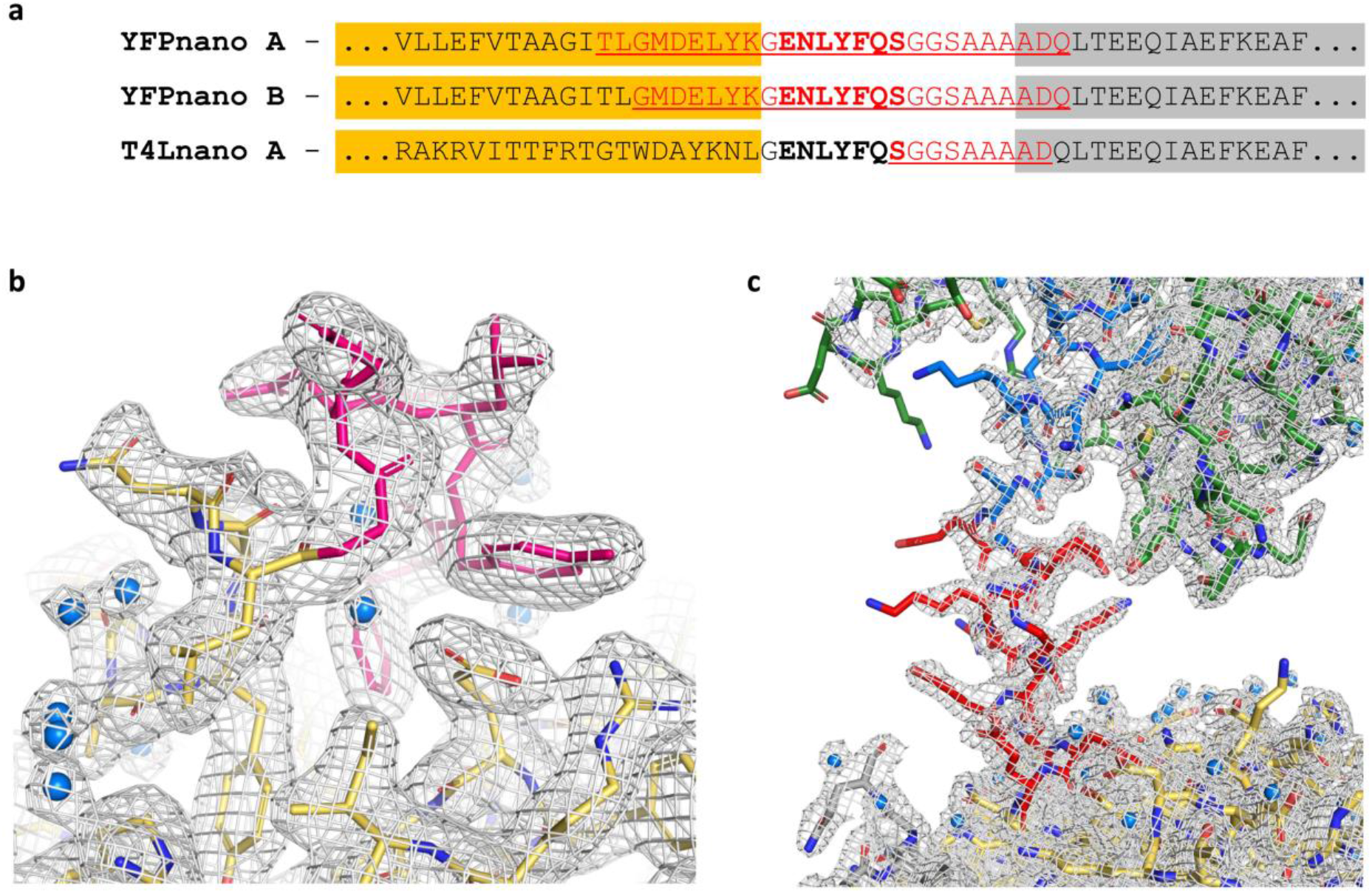
Crystal structure of the T4Lnano fusion protein - linker details. **a,** Sequence alignment of the region containing the flexible linker connecting YFP or T4L with CaM in the YFPnano/T4Lnano constructs. In the YFPnano crystal structure, there are two molecules in the asymmetric unit, the sequence of both chains (A and B) is shown here. YFP and T4L are highlighted in yellow. CaM is highlighted in grey. The TEV protease recognition site is shown in bold (the TEV site was not cleaved). Residues that were flexible and hence not visible in the crystal structures are shown in red font. In YFPnano, the flexible part of the linker counted >20 residues, allowing intramolecular CaM-to-CBP binding. In T4Lnano, the linker was designed shorter. Furthermore, the TEV site (bold) part of the linker engaged in well-ordered contacts with the T4L structure (b), resulting in further shortening of the linker. **b,** Close up view of the TEV site residues (pink) in the T4Lnano structure. T4L is shown in yellow and atom colors. Waters are depicted as blue spheres. 2Fo-Fc density map at a contour level of 1 σ displayed in a radius of 1.5 Å about each atom of the T4Lnano model. **c,** Close up view of the ER/K linker (red). The 2Fo-Fc density map is shown as in (b). CBP (blue) - ER/K motif (red) - T4L (yellow) - linker (not shown) - CaM (green).

**Supplementary Fig. 4).**
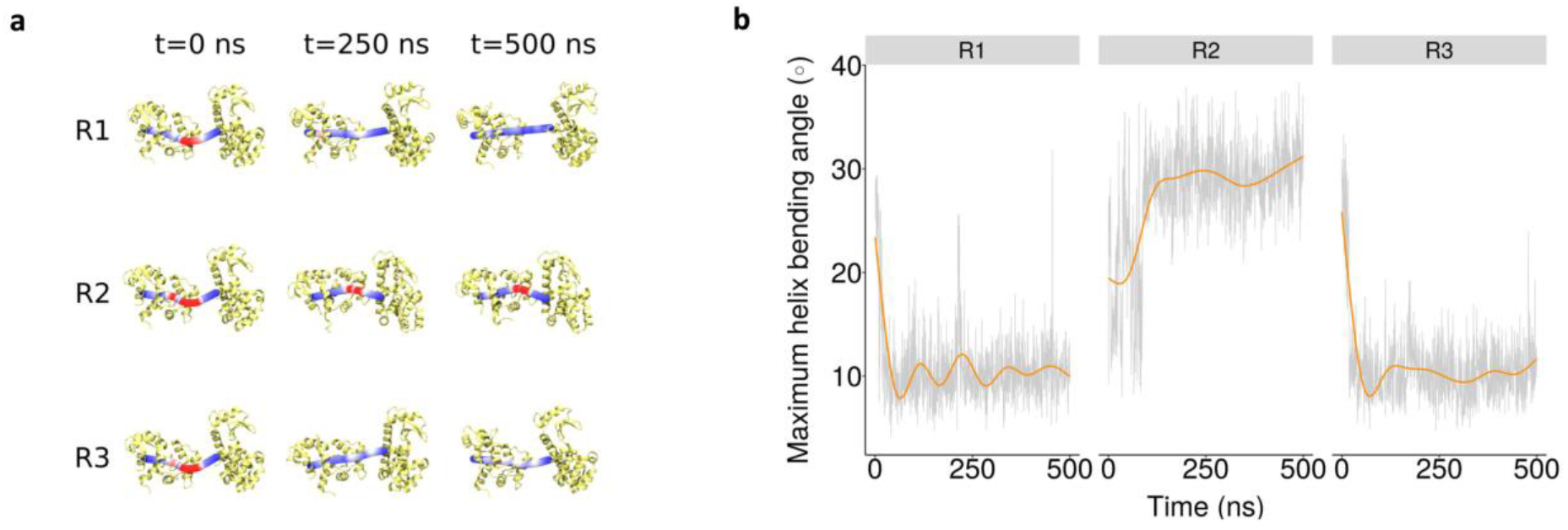
Extension of T4Lnano simulations. **a,** The panel shows three snapshots, namely at the beginning (0 ns), in the middle (250 ns), and at the end (500 ns), for each 3 x 500 ns simulation replicas of T4Lnano. Bendix^20^ was used to represent the curvature of the CBP-ER/K helix. A gradient color scale, namely blue (low) to red (high), was used to represent maximum angle values of this helix during the simulation. The rest of the protein is depicted in light yellow cartoons. **b,** Time series plots of maximum helix bending angles for the extended T4Lnano replicas. Maximum helix bending angles of the CBP-ER/K helix (i.e. residues 6 to 36) over time are depicted as light gray background lines across replicas (i.e. 3 x 500 ns). Overlaid bold orange lines are smoothed averages.

**Supplementary Movie 1)** Crystal contacts induce helix bending in T4Lnano - Representative simulation of T4Lnano comprising the first 100 ns of replica R1, where the CBP-ER/K helix (i.e. residues 6 to 36) clearly straighten up in the absence of crystal contacts. CBP (residues 6 to 25) and ER/K (residues 26 to 36) helices are represented in blue and yellow cartoons, respectively, and transparent surface. The rest of the protein is depicted using white cartoons. An overlaid red dashed line mimics a completely straightened helix.

## Supplementary Information on the CBP-CaM interaction

Overall, the CaM-CBP binding mode in our YFPnano and T4Lnano structures is in good agreement with the solution NMR structure described by Ikura et al.^17^, based on which our construct was designed. The calcium-sensing protein calmodulin consists of two domains **(Supplementary Fig.5)**.

**Supplementary Fig. 5).**
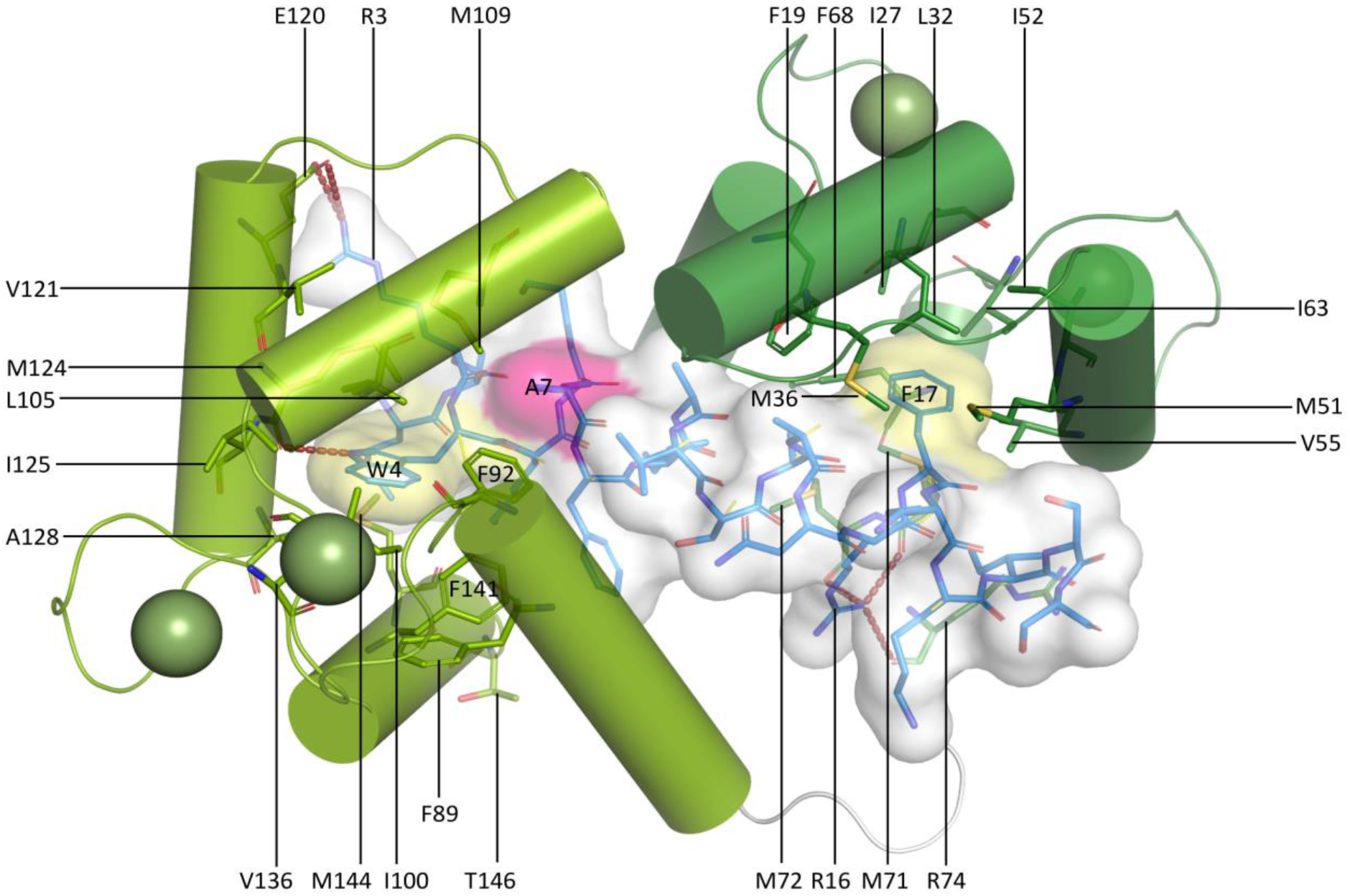
CaM - CBP interactions in T4Lnano. The calmodulin-binding peptide is shown as sticks (blue and atom colors) and as a transparent surface (white). Alanine 7, the affinity-increasing mutation, is highlighted in pink in the surface representation. Calmodulin is shown as a cartoon in green (light green = C-terminal domain, dark green = N-terminal domain). A selection of key residues is shown as sticks (CaM C = green, CBP C = blue, N = dark blue, O = red, S = yellow). The hydrophobic anchors W4 and F17 of the CBP are highlighted in yellow in the surface representation. Calcium ions are depicted as green spheres. Polar interactions are indicated by red dotted lines. The loop connecting the two CaM domains is shown in white.

The N-terminal domain comprises amino acids 4 −74 and the C-terminal domain comprises residues 82-147. The two domains are connected by a loop in the center (amino acids 75 - 81). Each domain encompasses two EF hand helix-loop-helix motifs, each coordinating one calcium ion. One CaM molecule can hence bind up to four calcium ions. Peptide binding by CaM is calcium-dependent^15, 18^. The two loops, each coordinating one calcium ion in one of the two EF hand motifs of a CaM domain, lie in an antiparallel direction to each other and engage in loop-to-loop, backbone-to-backbone hydrogen bonds between Ile 27 and Ile 63.

The sMLCK peptide is buried in a large hydrophobic channel of CaM, the CBP-CaM interaction is dominated by hydrophobic interactions. Tryptophan 4 and phenylalanine 17 of the CBP form hydrophobic anchors that insert into hydrophobic pockets that are located between the four helices of the two EF-hand motifs of the C-terminal, respectively of the N-terminal domain of CaM (**Supplementary Fig. 5** and Ref.^17^). CaM residues F92, I100, L105, M124, I125, A128, V136, F141, M144, M145, and more distantly F89, M109 and V121 provide a hydrophobic environment around CBP W4, while CBP F17 is embedded in a pocked that is formed by the CaM residues F19, I27, L32, M36, M51, I52, V55, I63, F68 and M71.

To assure formation of a very stable complex, we used a CBP sequence with a single-point amino acid mutation (Asn-to-Ala) in the MLCK M13 peptide that results in super-high affinity to CaM (K_d_ = 2.2(±1.4) × 10^−12^ M, corresponding to a ~1000-fold affinity improvement compared to the wt peptide)^15^. This residue is in proximity of the hydrophobic CaM residues F92, V108, M109 and L112 **(Supplementary Fig. 5)** and of the negatively charged amino acid E114. Alanine fits better into this mostly hydrophobic environment than the corresponding polar asparagine from the wild type peptide.

**Supplementary Table 1).** Data collection and refinement statistics

|  | YFPnano | T4Lnano | MoStoNano |
| --- | --- | --- | --- |
| Space group | $P2_1$ | $P2_12_12$ | $H32$ |
| Cell dimensions |  |  |  |
| $a, b, c$ (Å) | 53.7, 117.8, 84.5 | 86.38, 104.26, 63.95 | 165.69, 165.69, 352.16 |
| $\alpha, \beta, \gamma$ (°) | 90, 99.9, 90 | | |
| Resolution range (Å) | 48.59-1.901 (1.97-1.90) | 44.63-2.079 (2.15-2.08) | 47.84-2.85 (2.98-2.85) |
| $R_{\text{meas}}$ | 0.10 (1.02) | 0.13 (2.13) | 0.23 (1.14) |
| $CC_{1/2}$ | 0.998 (0.696) | 0.999 (0.6) | 0.998 (0.807) |
| $I/\sigma I$ | 14.12 (1.62) | 14.53 (1.31) | 8.2 (2.3) |
| Completeness (%) | 99.88 (99.57) | 99.84 (99.02) | 95.5 (99.7) |
| Multiplicity | 6.5 (5.1) | 13.1 (13.1) | 10.6 (10.3) |
| Total reflections | 526,754 (41,413) | 465,421 (44,996) | 351,924 (17,177) |
| Unique reflections | 81,149 (8,052) | 35,404 (3,434) | 33,351 (1,670) |
| $R_{\text{work}}/R_{\text{free}}$ | 0.1808 / 0.2138 | 0.1915 / 0.2358 | 0.2329 / 0.2625 |
| No. atoms |  |  |  |
| Macromolecules | 6,496 | 2,783 | 4,650 |
| Ligands | 124 | 37 | 63 |
| Solvent | 366 | 135 | 211 |
| B-factors |  |  |  |
| Macromolecules | 37.27 | 51.03 | 66.16 |
| Ligands | 39.98 | 64.49 | 81.81 |
| Solvent | 38.64 | 50.05 | 38.52 |
| R.m.s. deviations |  |  |  |
| Bond lengths (Å) | 0.009 | 0.011 | 0.014 |
| Bond angles (°) | 0.94 | 1.04 | 1.90 |

Highest-resolution shell in parentheses.

